# Neural activity tracking identity and confidence in social information

**DOI:** 10.1101/2021.07.06.449597

**Authors:** Nadescha Trudel, Matthew FS Rushworth, Marco K Wittmann

## Abstract

Humans learn about the environment either directly by interacting with it or indirectly by seeking information about it from social sources such as conspecifics. The degree of confidence in the information obtained through either route should determine the impact that it has on adapting and changing behaviour. We examined whether and how behavioural and neural computations differ during non-social learning as opposed to learning from social sources. Trial-wise confidence judgments about non-social and social information sources offered a window into this learning process. Despite matching exactly the statistical features of social and non-social conditions, confidence judgments were more accurate and less changeable when they were made about social as opposed to non-social information sources. In addition to subjective reports of confidence, differences were also apparent in the Bayesian estimates of participants’ subjective beliefs. Univariate activity in dorsomedial prefrontal cortex (dmPFC) and posterior temporo-parietal junction (pTPJ) more closely tracked confidence about social as opposed to non-social information sources. In addition, the multivariate patterns of activity in the same areas encoded identities of social information sources compared to non-social information sources.

## Introduction

To behave adaptively, humans and many other species learn from their conspecifics which actions to take and which ones to avoid. The human ability to learn not only from direct interactions with the environment, but also from other people contributes critically to our behavioural repertoires as individuals and also to our collective culture (Boyd and Richerson, 2009). When seeking information about which courses of actions to pursue, we routinely turn to others for advice and adjust our own course of action based on the quality of their advice (Bang et al., 2017, 2020; Behrens et al., 2008; De Martino et al., 2017).Clearly it is important to learn over time whether an advice giver is predicting external events accurately. This process, sampling and learning from other people’s advice, is arguably a process that may be very similar to learning from non-social cues about the occurrence of external events (Akaishi et al., 2016; Trudel et al., 2020). In other words, the cognitive operations underlying this type of information search might occur both in social and in non-social scenarios. Therefore, by comparing information sampling from social versus non-social sources, we address a long-standing question in cognitive neuroscience, the degree to which any neural process is specialized for, or particularly linked to, social as opposed to non-social cognition (Chang and Dal Monte, 2018; Diaconescu et al., 2017; Frith and Frith, 2010, 2012; Grabenhorst and Schultz, 2021; Lockwood et al., 2020; Soutschek et al., 2016; Wittmann et al., 2018). Given their similarities, it is expected that both types of learning will depend on common neural mechanisms. However, given the importance and ubiquity of social learning, it may also be that the neural mechanisms that support learning from social advice are at least partially specialized and distinct from those concerned with learning that is guided by non-social sources. Therefore, here, we seek to understand the behavioural and neural computations that differentiate learning from social and non-social information sources.

There is evidence that at least some parts of the brain are specialized for to negotiating social situations (Noonan et al., 2014; Sallet et al., 2011; Sliwa and Freiwald, 2017). For example, neuroimaging studies in macaques show a positive correlation between group size and areas in the temporal lobe (Sallet et al., 2011), in an area that appears to correspond to the temporoparietal junction (TPJ) in humans (Mars et al., 2013). In turn, TPJ has been associated and causally linked to the ability to infer other people’s beliefs (Hill et al., 2017; Schurz et al., 2017) and, in monkeys, to inferring the intended actions of others (Ong et al., 2021). Dorsomedial prefrontal cortex (dmPFC) is a second important node within this network again both in humans (Hampton et al., 2008; Nicolle et al., 2012; Piva et al., 2019; Suzuki et al., 2012; Wittmann et al., 2018) and in non-human primates (Noritake et al., 2018; Yoshida et al., 2012). For example, a recent study demonstrates dmPFC is causally implicated in social cognition and the maintenance of separate representations of oneself and of other individuals even when people interact (Wittmann et al., 2021). There are however, studies that report many common brain areas linked to reward and motivational processes in the context of both social and non-social behaviour suggesting at the very least that social information processing is not completely separate (Behrens et al., 2008; Boorman et al., 2013; Ruff and Fehr, 2014; Will et al., 2017; Zink et al., 2008). For example, tracking the probability of being correct or successful, often referred to as ‘confidence’ in one’s choice, has been shown to correlate with activation in perigenual anterior cingulate cortex (pgACC) in both social (Wittmann et al., 2016) and non-social (Bang and Fleming, 2018) settings.

It has been proven difficult to dissociate social as opposed to non-social cognitive functions. One reason is the lack of controlled within-subject experimental designs that have rarely been implemented in previous social neuroscience studies. Another reason might be the type of cognitive process that is studied; only cognitive functions that might plausibly occur in either social and non-social scenarios allow for a fair comparison. Here, we attempted to optimise our experimental design to address both these considerations in order to examine whether there are any fundamental differences in behavioural and neural computations when seeking information from social as opposed to non-social sources. First, we implemented a tightly controlled within-subject design with two experimental sessions that were matched in their statistical properties and only differed in their framing (social vs. non-social). This ensured that differences between conditions were confined to the nature of the information source -- social or non-social – and therefore that any differences in behaviour or neural activity could only attributed to the social/non-social difference. Second, we compared learning from other people’s advice with learning from non-social cues about the occurrence of external events. On every trial of the experiment, participants witnessed the precision with which a social cue (i.e., advice) or a non-social cue predicted a target. Over time, participants learned about the quality of these predictions and formed estimates of how confident they were in a cue, i.e., how precisely they expected the target to be predicted by the cue. We show that humans appear more certain in the performance accuracy of social predictors (advisors) compared to non-social predictors. This was evident in more stable confidence judgements across multiple timescales in the social as opposed to the non-social condition; there was a stronger reliance on representations made in the past and a weaker integration of new contradictory evidence. By using a computational approach, we could associate differences in the stability in confidence about social advisors to differences in the Bayesian estimates of participants’ subjective beliefs. We found that two brain areas, dmPFC and posterior TPJ (pTPJ) showed specificity in their processing of social information, not only in their average activation but also by encoding the identity of social cues in their multivariate pattern activation.

## Results

We used a social and non-social version of a previously validated paradigm (Trudel et al., 2020) during which participants learn about the performance of social and non-social cues in predicting a target on the circumference of a circle. On every trial, participants receive advice on where to find the target (Figure 1a,b, Supplementary Figure 1). In the social version, faces represented advisors that predicted a target position (a flower’s position on the circumference of a circle) (Figure 1a). Some advisors predicted the target accurately and others less accurately. Participants were instructed that advisors had previously been players of a similar task during which these players could directly learn about the target location (Figure 1c, Supplementary Figure 2). In other words, the other players had learned directly about the target location, while in the main experiment, participants could only infer the target locations from the predictions made by these previous players who now acted as advisors, but not directly from the target. Performance was defined by the size of an angular error, which represented the distance between the location of the predicted flower position (represented by a black dot) and the true position. In the non-social version, participants were instructed to collect fruits (this was now the target position and again it was represented by a yellow dot) that could either fall a small or large distance from the tree (this was now the predictor and again it was represented by a black dot). Importantly, task versions were completely matched in their experimental design and statistical properties inherent to the predictors and only differed in terms of their framing towards the participants. Thus, if there were differences in behaviour or neural activity between the conditions, then these could not have been attributed to any difference in protocol but only to a predisposition to treat the two types of predictors differently. On every trial, participants indicated their confidence in the advisor’s performance. A symmetrical confidence interval with a random initial width appeared around the (social or non-social) prediction. The confidence interval could be changed continuously to make it wider or narrower, by pressing buttons repeatedly (one button press resulted in a change of one step in the confidence interval). Reward was received when the target fell within the chosen interval and the number of points increased when the interval size was small. A narrow interval could be set when participants selected a predictor with a small angular error and when they were certain about the predictor’s angular error. Hence, trial-wise confidence judgments allowed us to inspect participants’ beliefs about the performance of the advisor and their uncertainty in this performance estimate. Beliefs about the predictor’s performance could be updated during the outcome phase through inspecting the angular error. Participants were presented with multiple social and non-social predictors across six blocks. In both social and non-social settings, predictors varied in the accuracy with which they predicted the target, i.e. their average angular error (Figure 1c). For each predictor, there was always another predictor in the other condition (social, non-social) with a matched accuracy level.

**Figure 1.**
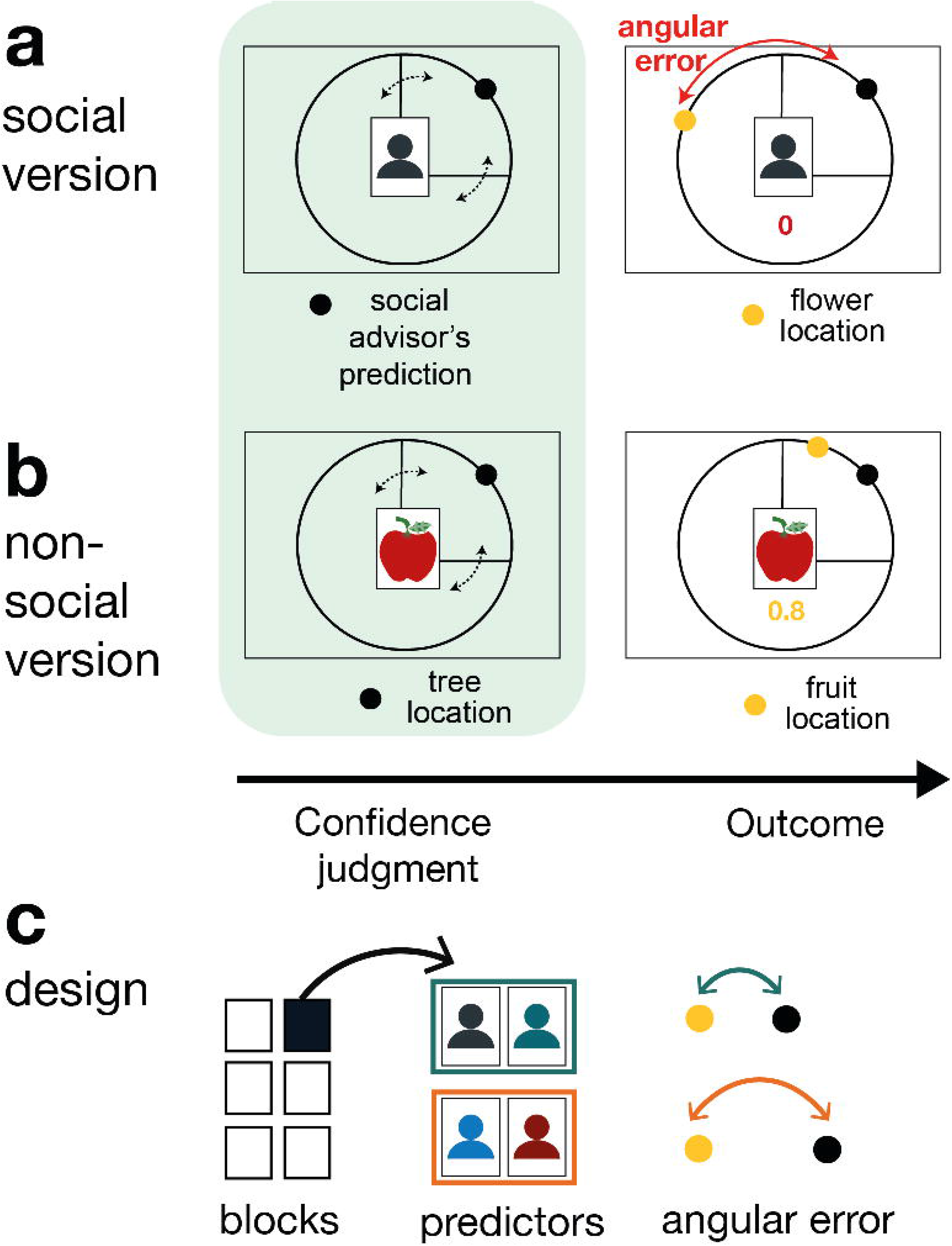
Social and non-social task versions and experimental design. Participants performed a social and non-social version of an information-seeking task. Versions only differed in their framing but were matched in their statistical properties. Participants learned about how well predictors (social advisors (a) and non-social cues (b)) estimated the location of a target on the circumference of a circle. On every trial, participants received advice on where to find the target. Participants indicated their confidence in the likely accuracy of performance by changing the size of a symmetrical interval around the predictor’s estimate (black dot, confidence phase): a small interval size indicated that participants expected the target to appear close to the predictor’s estimate. During the outcome phase, participants updated their beliefs about the predictor’s performance by inspecting the angular error, i.e., the distance between the predictor’s estimate (black dot) and true target location (yellow dot). **(a) Social version.** Information was given by social advisors that were shown as facial stimuli (note, that we are showing abstract persons in the manuscript, while facial stimuli were used in the experiment). Participants were instructed that advisors represented previous players that learnt about the target distribution themselves. Crucially, participants could not learn about the target (yellow dot) location themselves and could only infer the target locations from the predictions (i.e., social advice; black dot) from these previous players. **(b) Non-social version.** Participants selected between fruits (yellow dots) that fell with a different distance from their tree location (black dot) (note, that we are showing abstract fruits in the manuscript, while real fruit stimuli were used in the experiment). The aim was to select fruits with the smallest distance. Again, participants could not learn about the fruit location themselves, but had to infer it from the distance to their trees. (fruit stimuli depicted here are **(c) Design.** Each session comprised six experimental blocks (in total 180 trials). Each block included four new predictors (here as an example shown for the social condition with four abstract persons), of which there were two good predictors (on average small angular error) and two bad predictors (on average big angular error).

### Matching confidence to the performance of social and non-social advisors

We compared trial-by-trial confidence judgments between social and non-social predictors. When setting the interval, the aim was to minimize the discrepancy between the predictor’s angular error and the interval’s size and still include the target in the interval (Figure 2a, inset). Setting the interval in such a way allowed participants to maximize their reward across trials. Even though, on average, social and non-social sources did not differ in the precision with which they predicted the target (Supplementary Figure 3a), participants were better at adjusting their confidence judgment to the true performance of the social advisor compared to the non-social predictor. The confidence judgement was closer to the angular error in the social compared to the non-social sessions (Figure 2a, paired t-test: social vs. non-social, t(23)= −2.57, p= 0.017, 95% confidence interval (CI)= [−0.36 −0.4]).

**Figure 2.**
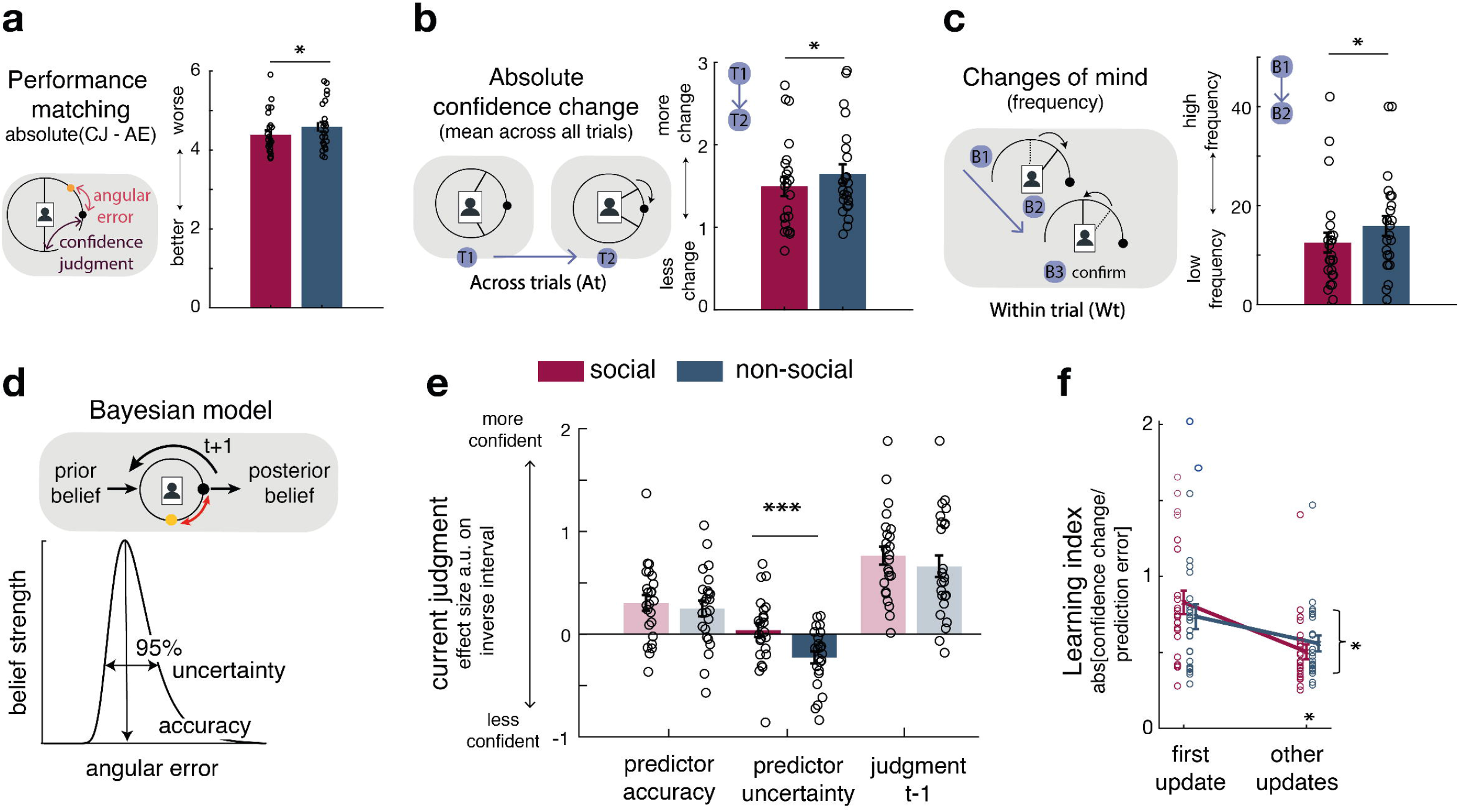
Confidence matching to the performance of social and non-social advisors. **(a)** Participants were more accurate in matching their confidence judgment (CJ) to the true performance (angular error; AE) of the social advisor (here represented as an abstract person rather than with a facial stimuli that was used in the experiment) compared to the non-social cue. **(b-c)** Changes of mind in confidence judgments are measured on two timescales: across trials, by comparing the absolute change in confidence judgments between the current and next judgment of the same predictor **(b)**; and within trials, by changes of mind just prior to committing to a final judgment **(c)**. Within trial changes of mind were defined by a specific sequence of the two last button presses (from button 1 (B1) to button 2 (B2)) before confirming the final judgment (button 3 (B3)): participants first decreased the interval (B1: more confident) and then increased it again (B2: less confident). Confidence judgments showed less changes of mind across trials (b) and within trials (c) for social advisors compared to non-social predictors. Note that panels a-c show raw scores. **(d) Bayesian model.** We used a Bayesian model to derive trial-wise belief distributions about a predictor’s performance. The prior distribution updates after observing the angular error of the selected predictor, resulting in a posterior distribution which serves as prior distribution on the next encounter of the same predictor. At each trial, two belief estimates were derived: first, the belief in the accuracy (point-estimate representing the maximum) and second, the uncertainty in that belief (95% range around the point-estimate) (more details, see Supplementary Figure 2). **(e) Confidence GLM.** We used a linear regression model to predict trial-wise confidence judgments (i.e., the width of the interval). Note that we have sign-reversed the confidence interval to make the measure more intuitive: a greater value corresponds to more confidence (i.e., smaller settings of the confidence interval). Participants were more confident when they selected a predictor they believed to be accurate. In addition, confidence judgments were impacted by subjective uncertainty in beliefs but only in the case of the non-social predictors: a larger interval was set when participants were more uncertain in their beliefs about the predictor’s accuracy. **(f) Learning index.** A trial-wise learning index shows that after the first update, changes in confidence judgments about social advisors are less strongly impacted by deviations between the current judgment and angular error (i.e., prediction error). (n=24, error bars denote s.e.m. across participants.)

Estimating the angular error of an advisor accurately depends on how judgments are adjusted from trial to trial. We compared changes in confidence judgments on two timescales. First, we did this on a longer timescale, by comparing the absolute change between the current and next judgment of the same predictor (which might only be presented again several trials later, Figure2b). Second, we did this on a within-trial timescale, by comparing changes of mind prior to the final judgment: participants adjusted the confidence interval’s size by individual button presses, allowing inspections of changes of mind within a trial according to a specific sequence of button presses prior to the final judgment. Changes of mind within a trial were indicated by first becoming more confident (selecting smaller intervals) and then becoming less confident (selecting a bigger interval) before the final judgment (Figure 2c, inset). Participants made fewer adjustments of their confidence judgements across both timescales for social advisors: participants’ absolute changes in their confidence judgments from trial to trial were lower (Figure 2b, paired t-test: social vs. non-social, t(23)= −2.7, p= 0.0125, 95% CI= [−0.3 −0.04]) and they exhibited fewer changes of mind before committing to a final judgment (paired t-test: social vs. non-social, t(23)= −2.4, p=0.025, 95% CI= [−6.3 −0.45]). In sum, these results suggest that participants are more certain in their interval choice for social compared to non-social information sources, as they change estimates about an advisor less drastically across trials and commit to their judgment within each trial with less hesitation.

We tested the hypothesis that participants’ certainty in their own beliefs influenced their confidence choice differently in social and non-social settings. We used a Bayesian model that allowed dissociation of two aspects of subjective beliefs: first, belief in how well the predictor will perform on a given trial (a belief in the predictor’s ‘accuracy’) and second, the participant’s uncertainty associated with the predictor’s predicted accuracy (‘uncertainty’ of the belief) (Figure 2d, Supplementary Figure 4) (Trudel et al., 2020). Both aspects of a belief might impact on the adjustment of the confidence judgment. Accuracy is the more likely belief parameter of the two to do so because accuracy reflects a point-estimate of the most likely angular error that an advisor has shown in the past. However, better performance matching (Figure 2a) can only be achieved when participants believe the advice given to be a true reflection of the target location (the advice is accurate) and when they are certain in their belief that the advice will be accurate (certainty in the advice’s accuracy). We tested the impact of both belief parameters on trial-wise interval settings and examined whether their impact differed when making judgments about social or non-social cues. For both social and non-social predictors, participants were more confident and set a smaller interval when they believed the predictor to be accurate in their target prediction (paired t-test: social vs. non-social, t(23)= 0.6, p=0.558, 95% CI= [−0.13 0.24]; one sample t-test: non-social, t(23)= 3.2, p=0.003, 95% CI= [0.09 0.41]; one sample t-test: social, t(23)=4, p<0.0001, 95% CI= [0.15 0.46], Figure 2e; Supplementary Figure 2). However, only when judging non-social predictors, confidence judgments were additionally widened as a function of subjective uncertainty: participants set a larger interval when they were more uncertain about the non-social cue’s accuracy to predict the target (paired t-test: social vs. non-social, t(23)= 3.5, p=0.0018, 95% CI= [0.11 0.42]; one sample t-test: non-social, t(23)= −3.9, p<0.0001, 95% CI= [−0.35 −0.1]; one sample t-test: social, t(23)=0.6, p=0.56, 95% CI= [−0.1 0.19], Figure 2e). Uncertainty about the accuracy of social advisors, however, did not have the same impact: participants’ confidence in the social advisors were instead guided directly by the peak of the Bayesian probability distribution, as indicated by the accuracy effect.

We have seen that participants have less changeable estimates of the performances of social advisors compared to non-social advisors (Figure 2b), and this relative increase in certainty about the value of social advice as opposed to non-social information is also reflected in the Bayesian estimates of participants’ subjective beliefs (Figure 2d,e). One possibility is that this increased certainty is caused by relying on a longer-term memory of the observed advice in the social compared to the non-social context. Participants might form an opinion about the performance of social advisors more quickly and are therefore less likely to change it in face of new and possibly contradictory evidence. We therefore tested whether the degree of error-driven behavioural adaptation is different between social and non-social conditions. In other words, we tested whether participants changed their judgment from one trial to the next to a smaller degree as a function of the discrepancy between their judgments and the angular error (i.e. prediction error). We calculated a trial-by-trial learning index, similar to a learning rate in a reinforcement learning model (Sutton and Barto, 1998; Wittmann et al., 2020), that measured the change between the current and next judgment given the prediction error (Figure 2f; Methods, Equation 2, 3). A smaller learning index shows less error-driven adjustment of one’s confidence judgment. Learning indices between conditions were similar for the first observation but declined more steeply for social compared to non-social predictors for the remaining updates (repeated-measures ANOVA: interaction between group (social, non-social) x timepoint (first update, other remaining updates), F(1,23)= 5.8, p=0.024, Figure 2f). Hence, after the first update, social judgments were subsequently less influenced by prediction errors that were observed (paired t-test: social vs. non-social for remaining only, t(23)= 2.7, p=0.01, 95% CI= [−0.95 −0.13], Figure 3f). As a consequence, judgements relied more on observations made further in the past. This is consistent with the observation that participants repeatedly set similar confidence intervals across time in the social condition (i.e. fewer changes across trials) (Figure 2b), and that these settings were less impacted by their own subjective uncertainty (Figure 2e).

**Figure 3.**
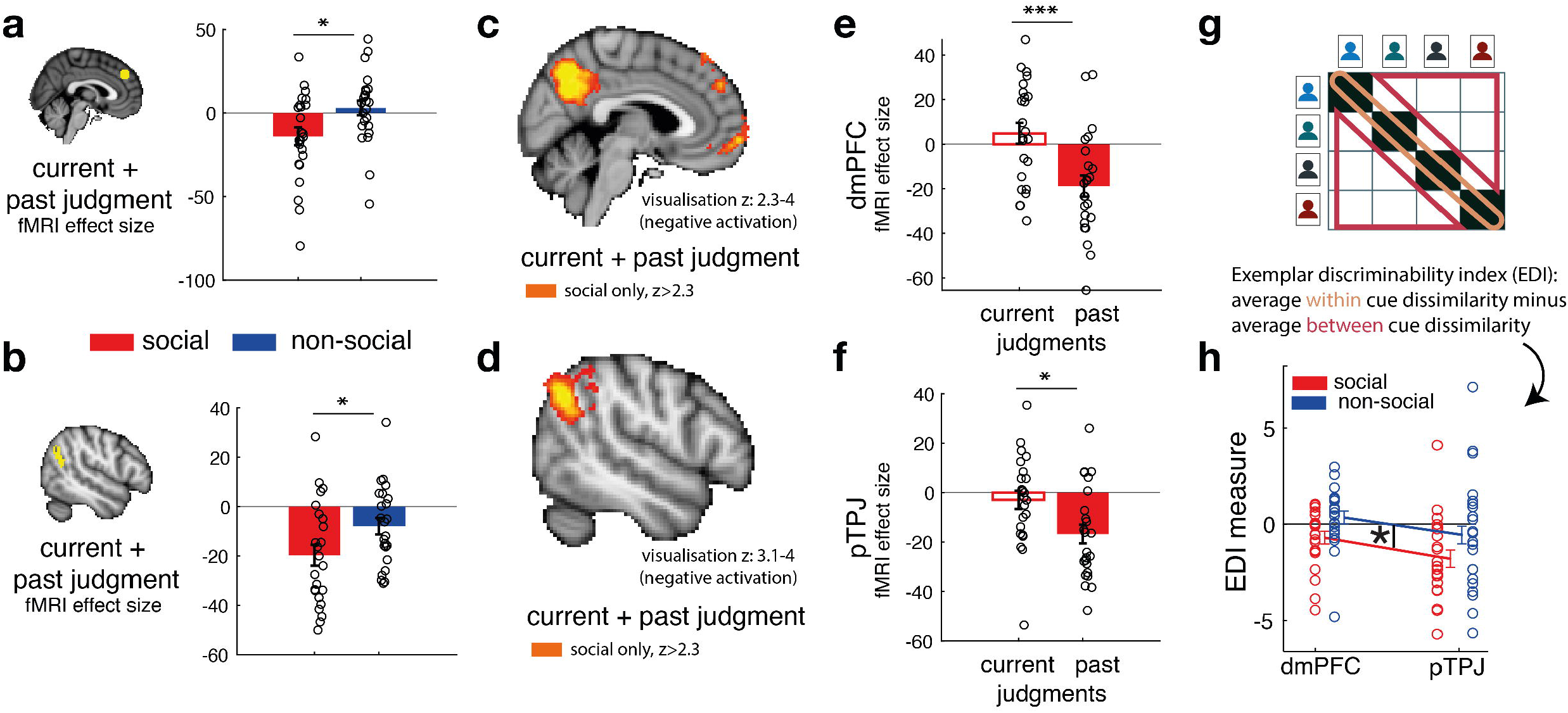
Social confidence and advisor identity in dmPFC and pTPJ. We used a region-of-interest approach based on two independent *a priori* regions in dmPFC **(a, left)** and pTPJ **(b, left).** Activation in both areas covary with the combination of current and past judgments for social advisors compared to non-social advisors. We performed a whole-brain exploratory analysis and show cluster-corrected activation for the same contrast as in (a,b) for the social condition in dmPFC **(c)** and pTPJ **(d)**. Analyses were FWE cluster corrected with z-score> 2.3 and p<0.05, but for visualisation purposes, z-thresholds differ between both images (dmPFC, z: 2.3-4 and pTPJ, z: 3.1-4). Note that we use the same color scheme to indicate social (red color) and non-social conditions (blue color) throughout this figure; however, the depicted whole-brain activations are of negative polarity (Supplementary table 1). Both dmPFC **(e)** and pTPJ **(f)** represent judgments that are informed by past observations compared to current observations (see Supplementary Figure 5 for whole-brain results) in the social condition. Activation in both areas scaled significantly more strongly with past compared to current judgments for social advisors. **(g)** A representational similarity analysis was applied to the BOLD activity patterns evoked by each predictor (faces and fruits in the social and non-social conditions, respectively; note that we are using abstract persons in the manuscript while we used real faces in the experiment) at the time of response during the confidence phase. We compared pattern dissimilarity across voxels (measured by the Euclidean distance) between all pairwise combinations of social and non-social cues. Exemplar discriminability index (EDI) was calculated to compare the dissimilarity of the same cues (across diagonal, orange area) to the dissimilarity of different cues (off-diagonal, red area). **(h)** A negative EDI indicates multivariate activity pattens are more similar when the same cue, as opposed to different cues are seen. Multivariate patterns within dmPFC and pTPJ encoded the identity of social cues more strongly than it encoded the identities of non-social predictors. (n=24, error bars denote s.e.m. across participants; whole brain-effects are family-wise error cluster corrected with z-score> 2.3 and p<0.05).

### dmPFC and pTPJ covary with social confidence and encode social advisor identity

Next, we tested whether there are brain areas that covary with confidence judgments (i.e. the width of the confidence interval set by the participants) that are made about social compared to non-social advisors. Behaviourally, we have shown that judgments about social advisors might be particularly strongly impacted by relatively constant and unchanging representations held in memory. Therefore, a combination of the trial-wise confidence judgments made in the current and past trials for the same predictor was used to test for differences between social and non-social advisor judgments. We sign-reversed the relationship of the confidence measure such that a higher confidence index now represented greater confidence in the selected predictor. We focused on two *a priori* independent ROIs, dmPFC (Wittmann et al., 2016) and pTPJ (Mars et al., 2013) because of their previous association with inference about beliefs held by others (Hampton et al., 2008; Nicolle et al., 2012; Piva et al., 2019; Saxe, 2006; Schurz et al., 2017; Suzuki et al., 2012; Wittmann et al., 2016); a process that is crucial to the current task. Hence, we tested whether blood-oxygen-level dependent (BOLD) signal in these brain areas changed in tandem with variation of the expressed confidence. This parametric contrast captures trial-by-trial variation in the confidence that participants have in the predictor and is independent of the mean confidence levels. Note that the mean confidence level was not significantly different between social advisors and non-social cues (paired t-test: mean confidence, social vs. non-social, t(23)= −1.7, p=0.11, 95% CI= [−0.66 0.07]), even though we observed greater stability of confidence for social advisors (Figure 2b). Activation in both dmPFC and pTPJ covaried with combined confidence judgments significantly more strongly for social compared to non-social advisors (Figure 3a, paired t-test: social vs non-social for dmPFC, t(23)= −2.5, p=0.019, 95% CI= [3 31]; Figure 3b, paired t-test: social vs non-social for pTPJ: t(23)= −2.3, p=0.03, 95% CI= [1 22.5]). Next, we performed an exploratory whole-brain analysis and found that these activations in dmPFC and pTPJ were further cluster-corrected when making judgments about social advisors (family-wise error (FWE) cluster corrected with z-score> 2.3 and p<0.05, Supplementary table 1). In addition, activity was whole-brain cluster-corrected in some other areas associated with social cognition such as the medial frontal pole, posterior cingulate/medial parietal cortex for confidence judgments about social advisors (Figure 3c,d, for a complete list of cluster-corrected brain areas, see Supplementary table 1).

Our behavioural results showed that social judgments might be especially impacted by beliefs formed over longer timescales: for example, judgments about social advisors changed less drastically across time, even if new information at the time of the trial outcome was contradictory to one’s current belief (Figure 2f). We tested whether such a differentiation between current and past confidence judgments was also apparent in the neural activation profile of dmPFC and pTPJ. In other words, we tested whether pTPJ and dmPFC displayed an activation profile consistent with representing past judgments about social advisors more strongly in relation to current judgments about the same advisors. Consistent with behavioural results, activation within both areas covaried with judgments made on longer time-scales more strongly than present judgments (one sample t-test, social: dmPFC, past vs current: t(23)= 3.04, p= 0.0058, 95% CI= [7.6 39.8]; pTPJ, past vs. current, t(23)= 2.25, p= 0.034, 95% CI= [1.1 26.6]; Figure 3e,f; Supplementary Figure 5a). There was no difference in neural activity for confidence judgments between both of these timescales for non-social cues (Supplementary Figure 6).

So far, we have focussed on how the univariate activity in dmPFC and pTPJ covaried with the participants’ estimates of the accuracy of predictors. Next, we examined whether the same brain regions carried additional information about the predictors that is orthogonal to what we tested so far; we sought evidence as to whether they encoded the identities of the cues. We applied a representational similarity analysis (RSA) to BOLD activity patterns evoked by each face and each fruit presentation at the time of response during the confidence phase. We compared the pattern dissimilarity across voxels (measured by the Euclidean distance) in the same *a priori* independent ROIs in dmPFC and pTPJ as examined previously. For each condition, the Exemplar Discriminability Index (EDI) was calculated which represents the average dissimilarity within the same cue (Figure 3g, orange area) compared to the average dissimilarity between different cues (Figure 3g, red area) (Bang et al., 2020). Hence, a negative EDI indicates that pattern activations encode representations of the same cues compared to dissimilar cues. Pattern activations in both dmPFC and pTPJ encoded the identities of social cues more strongly than non-social cues (Figure 3h; repeated-measures ANOVA: main effect of condition (social, non-social) in 2 (condition: social, non-social) x 2 (area: pTPJ, dmPFC) analysis of variance: F(1,21)= 5, p= 0.036). In summary, in conjunction, the univariate and multivariate analyses demonstrate that dmPFC and pTPJ represent beliefs about social advisors that develop over a longer timescale and encode the identities of the social advisors.

## Discussion

We tested the behavioural and neural computations that differentiate learning from social and non-social information sources. Using a tightly controlled within-subject design in which conditions only differed in their social or non-social framing, we asked participants to make trial-wise confidence judgments about the advice of social and non-social information sources. Ideally confidence increases with the accuracy of one’s choice (Bang and Fleming, 2018; Kiani and Shadlen, 2009; Lak et al., 2020). Here, participants did not match their confidence to the likely accuracy of their own performance, but instead to the performance of another social or non-social advisor. Participants were better at matching their confidence judgments to the performances of social compared to non-social advisors. A possible explanation might be that participants have a better insight into the abilities of social cues – typically other agents – than non-social cues – typically inanimate objects. We used a computational approach to test the interplay of two belief estimates that impact confidence judgments: the belief in how accurate the advice is and the subjective uncertainty in that belief. Confidence judgments were impacted by a multiplicity of variables; however, it was the subjective uncertainty about the performance estimates that dissociated judgments about social and non-social advisors. Participants selected a greater interval size when making their confidence judgements on non-social trials, indicating that they were more uncertain about the performance accuracies of the non-social cues compared to the social cues.

Being less influenced by ones’ uncertainty about the value of social advice might help explain why people sometimes fail to use evidence to update their assessments of social advisors (Fleming et al., 2018; Rollwage et al., 2020). Detecting uncertainty in environmental contingencies or about one’s own beliefs is achieved with fine-tuned metacognitive abilities (Miyamoto et al., 2021) and is essential for behavioural adaptation (Badre et al., 2012; Behrens et al., 2007; Trudel et al., 2020). Participants were less influenced by their subjective uncertainty when estimating the performances of social advisors. Moreover, they changed and updated their beliefs about social cues less than about non-social cues when they observed the outcomes of their predictions. Repeating a particular interval size instead of adjusting it from trial to trial might be a better strategy in the current experiment because all advisors had, on average, a stable performance level. However, opting for such a strategy only in the context of social advisors might reflect a natural tendency to expect other social agents to behave consistently and exhibit stable performance levels as soon as an initial learning period is passed. It may be because we make just such assumptions that past observations are used to predict performance levels that they are likely to exhibit next (Wittmann et al., 2021, 2016). An alternative explanation might be that participants experience a steeper decline of subjective uncertainty in their beliefs about the accuracy of social advice resulting in a narrower prior distribution during the next encounter of the same advisor. From a Bayesian perspective, greater certainty about the value of advice means that contradictory evidence will need to be stronger to alter one’s beliefs. In the absence of such evidence, a Bayesian agent is more likely to repeat previous judgments. Just as in a confirmation bias (Kappes et al., 2020), such a perspective suggests that once we are more certain about others such as their traits, we are less likely to change our opinions about them.

We have shown that there are at least some unique processes involved when judging one’s confidence about social advisors compared to non-social cues. It may also be that the neural mechanisms that support learning from social advice are at least partially specialized and distinct from those concerned with learning that is guided by non-social sources. In particular two brain areas, dmPFC and pTPJ, have not only been shown to carry signals associated with belief inferences about others but, in addition, recent combined fMRI-TMS studies have demonstrated the causal importance of these activity patterns for the inference process (Hill et al., 2017; Wittmann et al., 2021). Similar processes might be at play in the current study during which participants make inferences about the performance of the social advisor in locating the target. We examined dmPFC and pTPJ, using ROIs derived from previous studies, and showed that these areas represent a combination of present and past confidence judgments that are made about social advisors and that they do this more strongly for social than for non-social cues. A long-standing question in cognitive neuroscience is the degree to which any neural process is specialized for, or particularly linked to, social as opposed to non-social cognition (Chang and Dal Monte, 2018; Diaconescu et al., 2017; Frith and Frith, 2010; Grabenhorst and Schultz, 2021; Lockwood et al., 2018, 2020; Ruff and Fehr, 2014; Rushworth et al., 2012; Soutschek et al., 2016; Wittmann et al., 2018). The current results suggest that dmPFC and pTPJ activity patterns are different during social cognition and that they carry information that is related to the performance differences that are seen at a behavioural level during a social as opposed to non-social task. Interestingly, it seems that activation in dmPFC and pTPJ was particularly prominent when representing judgments from previous encounters compared to the present judgment. This was apparent not only in univariate activity in these areas that covaried with confidence in advice but also in the multivariate activity patterns that encoded the identity of social agents; representational similarity analysis showed that pattern activations in both areas encoded the identity of social cues compared to the identity of non-social cues. Hence, pTPJ and dmPFC seem particularly linked to the navigation of social interactions.

## Methods

### Participants

30 participants took part in the experiment of which some were excluded because they fell asleep during the scan (N=2), exhibited excessive head motions (N=1) or because they prematurely terminated the sessions (N=3). The final sample consisted of 24 participants (14 female, age range 19-35). Each participant took part in two versions of the experiment: a social and non-social version. The order of versions was counterbalanced across participants. No statistical method was used to pre-determine the sample size. However, the sample size used is larger than those reported in previous publications (Boorman et al., 2011). Moreover, we are using a within-subject design to increase statistical power to detect a true difference between conditions. The study was approved by the Central Research Ethics Committee (MSD-IDREC-C1-2013-13) at the University of Oxford. All participants gave informed consent.

### Experimental procedure

All participants performed two sessions on separate days that only differed in their framing of the experimental task (social or non-social). Each session consisted of one-hour of magnetic resonance imaging (MRI) scanning and 30 minutes of instruction including practice trials outside the scanner. Participants learnt the trial procedure and response mapping during practice trials. Practice trials were independent from the main task; they included different stimuli and data and thereby guaranteed that participants encountered predictors for the first time during the scanning session. Participants received £15 per hour and an additional bonus depending on task performance (per session: £5-£7). After both sessions were completed, participants completed a debriefing questionnaire probing their understanding of the task. We used data previously reported by Trudel and colleagues (2020) but now with a fundamentally different aim. Previously Trudel and colleagues had focussed on uncertainty guided exploration of reward-predicting cues regardless of whether the cues were social or non-social in nature. Accordingly, their analyses collapsed across data obtained in the social and non-social conditions. Now, the question of exploration was no longer of primary importance but instead the focus was on the impact of social versus non-social cue differences on behaviour and neural activity. Here we investigated behaviour and neural activity in the second phase of each trial during which participants reported their confidence judgments. By contrast Trudel and colleagues (2020) focussed on behaviour and neural activity in the first phase of each trial, the decision phase (Supplementary Figure 1).

### Framing of social and non-social task versions

Each participant performed a social and non-social version of the task (Figure 1). Tasks were matched in their statistical properties, and only differed in the stimuli (predictors that were used to indicate an information source). In the social task, the predictors were represented with facial stimuli (stimuli IDs were as follows: AF02NES, AF06NES, AF06NES, AF07NES, AF08NES, AF11NES, AF19NES, AF23NES, AF24NES, AF25NES, AF27NES, AF30NES, AF32NES, AM04NES, AM08NES, AM10NES, AM11NES, AM14NES, AM22NES, AM23NES, AM25NES, AM26NES, AM27NES, AM28NES, AM35NES, (Lundqvist et al., 1998)). Note that in all figures we show abstract persons, while in the experiment participants saw facial stimuli of real people. They represented advisors who predicted a target position (a flower’s position on the circumference of a circle represented by a yellow dot). Advisors differed in in how well they predicted the target location and the participant’s aim was to select advisors who made target predictions that were closest to the subsequent actual target positions; if participants followed the guidance of more accurate predictors then they were more likely to win more points (see *Experimental design* section below) which ultimately led to a higher payout. In the non-social version, participants were instructed that they were to collect fruits (Foroni et al., 2013) that again lay on the circumference of a circle and which were again represented by yellow dots). Different fruits fell with different distances from their tree and the aim was to select the fruits that fell the smallest distance from their trees. Thus, the fruit trees served an analogous role to the faces in the social condition. In both cases the stimuli were predictors that estimated the subsequent position of yellow dotes on the circumference of a circle. Just as the advisors in the social condition varied in the accuracy with which they made their predictions about the subsequent yellow dot position, the fruit trees also varied in the accuracy with which they made their predictions about the subsequent yellow dot position. Crucially the statistical relationships between predictors and predicted items were varied in identical ways across both the social and non-social stimuli. In both conditions there were some more accurate and less accurate predictors (discussed more in Experimental design). Thus, if there were differences in behaviour or neural activity between the condition then this could not have been attributed to any difference in protocol but only to a predisposition to treat the two types of predictors differently. Stimuli for both task versions were drawn from a database and randomised across participants.

During the experimental task, participants could not learn about the target location directly, but instead had to make a binary selection between predictors during the first phase of the trial, the decision phase (Supplementary Figure 1). Having selected a predictor, participants received information on where to find the target in the second phase of the trial (confidence phase). In the social version, participants were instructed that the predictors (advisors in this case) represented previous players who had already performed a behavioural task during which they had had the opportunity to learn about the distribution of the target’s location (Supplementary Figure 2). During this prior behavioural pre-task, participants were told that the other players had learnt to predict the location of the target through trial and error across time. In other words, during the pre-task, the other players had learned directly about the target location, while in the main experiment, participants could only infer the target locations from the predictions made by these previous players who now acted advisors, but not directly from the target. Analogously in the non-social version of the task, participants were told that different fruits were dispersed more or less closely to their trees, so the tree position was a similar more or less accurate predictor of the target position. The deviation between the predictor’s target prediction and the true target position defined the angular error. It was those angular errors that were presented to participants in the main experiment and that enabled participants, albeit indirectly, to estimate the target location. Therefore, in the main experiment, participants were incentivized to try and identify predictors that were more accurate in their predictions and participants were able to do this by observing the angular errors associated with their predictions (Figure 1A). This constituted the only way for the participant to estimate the target’s location in the main experiment.

### Experimental design

In the experimental sessions, the participants’ aim was to select predictors that best predicted the target’s location on the circumference of the circle. Participants could learn about a predictor’s performance by selecting it in the first phase of the trial (decision phase) and observing the difference between the target prediction and the subsequent true target location (outcome phase) – we refer to this as the angular error (Figure 1 and Supplementary Figure 1). In the second phase of the trial, participants adjusted the size of a symmetrical interval, on a computer monitor, around the predictor’s prediction. The interval could be changed by individual button presses that either increased (to indicate less confidence) or decreased (to indicate more confidence) the interval. A third button was used to confirm the final judgment. The interval changed with a precision of 20 steps on each side of the predictor’s estimate. The aim of the participants was to select a predictor that predicted the target well. This in turn allowed participants to select a narrow confidence interval around the prediction. When this confidence interval included the target (as revealed in the subsequent outcome phase), then this resulted in a positive reward outcome. The positive outcome was higher if the confidence interval was set very narrowly, and the target still fell within it. If the target fell inside the interval, the payoff magnitude was determined by subtracting the interval size from 1. If the target fell outside the interval, then participants received a null payoff. Therefore, the payoff ranged between 0 and 1 as follows:

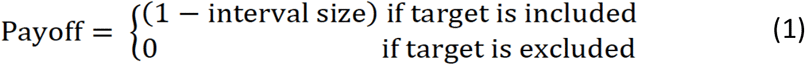

Participants transitioned through six blocks, during which they encountered four predictors each depicted by new stimuli in each block. The session comprised two blocks of 15, 30 and 45 trials each. Therefore, both social and non-social sessions comprised 180 trials each. The number of trials remaining was explicitly cued on every trial in the task (Supplementary Figure 1). In each part of the experiment (social and non-social), two predictors were associated with a better performance. They had on average a smaller angular error compared to the other two predictors in the block. The predictors’ estimates were determined by a Gaussian distribution centred on the true target location but with different standard deviations (standard deviation was either 50 or 70 for the better and worse predictors, respectively). A smaller standard deviation resulted in smaller angular errors. Hence, averaging across the angular errors they had observed allowed participants to learn about the standard deviation associated with a predictor’s true performance distribution. For more details, see Supplementary Figure 3.

### Bayesian model

We used a Bayesian model (Trudel et al., 2020) (Supplementary Figure 4) to guide our analyses. We focussed, first, on how participant’s behaviour might differ between the social and non-social situations and, second, on how participants estimated their uncertainty about the predictions which was revealed by their setting of the confidence interval. We submitted the computational variables from our model to a general linear model (GLM) analysis to predict confidence judgments. For every trial, we extracted two model-based subjective belief parameters that captured two key features of each participant’s likely beliefs about a predictor’s performance. First, the belief of how accurately a predictor would predict the target was captured by the mode of the probability density function on each trial associated with the selected predictor. Second, the subjective uncertainty in that accuracy belief was determined by a 95% interval around the mode. See Supplementary Figure 4 for more details on the Bayesian model.

### Behavioural analyses

To test for differences between social and non-social conditions, we first compared how participants set their interval during the confidence phase. We tested whether there were differences in how they learnt about a social versus a non-social predictor’s performance. We compared their performance in matching their confidence judgment with the true angular error. We computed an index for “performance matching” defined as the absolute difference between the true angular error and the size set for the confidence interval. We averaged the resulting indices over sessions and compared the performance matching index in the social and non-social sessions. Next, we tested how participants adjusted their confidence judgments across two timescales. First, we tested confidence adjustments across trials by calculating – for the same predictor - the absolute difference between the current and next size of the confidence interval averaged across all trials within social and non-social conditions (Figure 2b). Next, we exploited the fact that the interval size was modified by individual button presses, which allowed testing for “changes of mind” within a trial. Changes of mind were defined by a specific button sequence that first decreased (more confident) the interval and then increased (less confident) the interval prior to committing to the final judgment. We identified the number of trials per session that were characterised by such a “decrease – increase – commit” sequence and compared their frequency between social and non-social conditions (Figure 2c).

Next, we used a computational approach to disentangle observed differences in confidence judgments across conditions. We applied a general linear model (GLM) to predict changes in confidence judgments as a function of model-based and model-free regressors. Note that confidence judgments were modified by changing the interval size, with a smaller interval size indicating higher confidence. To make this measure more intuitive, we sign-reversed the relationship such that a higher confidence index now represented greater confidence in the selected predictor. Regressors were normalised across all trials (mean of zero and standard deviation of one). The GLM applied to the confidence phase comprised the following regressors and relevant effects are shown in Figure 3A (whole GLM is shown in Supplementary Figure 3): predictor accuracy, predictor uncertainty, payoff t-1 (payoff observed on the last trial (t)) for the same predictor, confidence judgments on the past three consecutive trials (t-1, t-2, t-3) for the same predictor and the initial starting position of the confidence interval were added (the latter represents a confound regressor).

Next, we tested the degree to which participants changed their confidence interval from one trial to the next according to the difference between the current interval that the participant had set when making their estimation of target position and the actual angular error that was subsequently observed. This difference can be thought of as the participant’s prediction error in this task. We calculated a learning index that resembles a learning rate in a reinforcement learning (RL) model, capturing the rate at which evidence observed at the time of the prediction error is integrated into the next estimate. Note that RL models are often fitted to the data to derive a trial-by-trial value estimate, to predict choice behaviour and neural correlates although direct observation of the model parameters is not possible. However here we are provided, by the participants, with explicit confidence judgments, allowing us to derive a learning rate index for every trial. We derived a trial-by-trial learning index by dividing the absolute change in confidence interval (CI) that participants set from one trial, t, to the next, t+1, by the unsigned prediction error (i.e., absolute difference between the CI on trial t and the angular error (AE) on trial t):

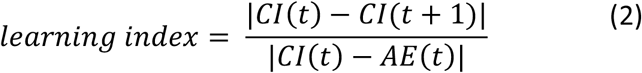

The magnitude of the learning index can be interpreted as follows: Zero indicates that participants did not change their confidence interval from one trial to the next. 1 indicates a confidence judgment change from one trial to the next that that is equivalent to the distance that their confidence interval was away from the target position in the last trial, which would be beneficial if the angular error for the predictor stayed the same across trials. Values bigger than 1 indicate over-adjustment: a change in confidence interval that is even bigger than the prediction error the participant had observed (this would happen if participants were expecting the angular error to be even greater on the next trial). Values smaller than 1 indicate under adjustment; participants are updating their confidence intervals by less than the prediction error that they observed.

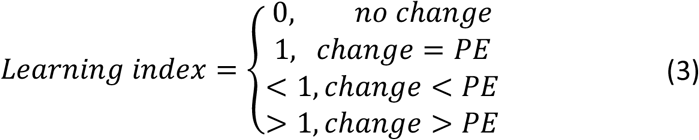

### Imaging data acquisition and pre-processing

Imaging data was acquired with a Siemens Prisma 3T MRI using multiband T2*-weighted echo planar imaging sequence with acceleration factor of two and a 32-channel head-coil(Trudel et al., 2020). We acquired slices with an oblique angle of 30 degrees from posterior to anterior commissure to reduce frontal pole signal dropout. The acquisition parameters were: 2.4×2.4×2.4mm voxel size, echo time (TE)=20ms, repetition time (TR) = 1.03ms. A fieldmap was acquired for each session to reduce distortions and bias correction was applied directly to the scan. The structural scan had a slice thickness of 1mm, TR =1.9ms, TE=3.97ms, and a voxel size of 1×1×1mm.

We used FMRIB Software Library (FSL) to analyse imaging data (Smith et al., 2004). Pre-processing stages included motion correction, spatial distortion correction by applying the fieldmap, brain extraction, high-pass filtering and spatial smoothing using full-width at half-maximum of 5mm. Images were first registered to each participant’s high resolution structural image and then non-linearly registered to the Montreal Neurological Institute (MNI) template using 12 degrees of freedom.

### Region-of-interest (ROI) approach

We used a ROI approach to test differences between social and non-social conditions in two regions. We extracted independent ROIs from dmPFC and pTPJ. dmPFC ROI was calculated with a radius of three voxels centred on MNI-coordinate [x/y/z: 2,44,36] (Wittmann et al., 2016). pTPJ ROI consisted of a 70% probability anatomical mask including both left and right hemispheres. We derived the left pTPJ ROI by warping the mirror-image of the right pTPJ (Mars et al., 2013) to the left hemisphere. The selected ROIs were transformed from MNI space to subject space and relevant COPEs (contrast of parameter estimate) were extracted for each participant’s session (Figure 3a,b,e,f).

### Exploratory MRI whole-brain analysis

We used a single GLM to analyse the data. All phases (decision, confidence and feedback phase) were included into the GLM. The decision phase was time-locked to the onset of predictor display. The confidence phase was time-locked to the response made in the confidence phase. The feedback phase was time-locked to the onset of outcome. The duration between trials was drawn from a Poisson distribution with the range of 4s to 10s and a mean of 4.5s. To decorrelate variables of interest between trial phases, short intervals were included between trials (inter-trial-intervals, ITIs) and randomly, but equally allocated to either the transition between decision- and confidence phase or confidence- and outcome phase. Each phase was modelled as constant. We included additional normalised parametric regressors (mean of zero and standard deviation of one) that were modelled as stick functions (i.e. duration of zero). The following parametric regressors were included:

#### Decision phase

- chosen-unchosen uncertainty
- chosen-unchosen accuracy.

#### Confidence phase

- current confidence judgment (i.e., the interval size selected on the current trial),
- past confidence judgment (i.e., the interval size selected on the previous encounter of the same predictor),
- number of button responses (that were made to reach the desired interval size; this regressor acted as confound regressor).

#### Feedback phase

- payoff.

We calculated the following contrasts of for the confidence phase: (current + past confidence judgment) and (current – past confidence judgment). In addition, we included one regressor that was time-locked to all button responses. Note that we only analysed the trials that provided both current and past judgments for the same predictor. The results were submitted to two second-level analyses applied separately to social and non-social conditions using FLAME1 (Supplementary Figure 5). All results were FWE cluster-corrected at p<0.05 using a cluster-defining threshold of z>2.3.

### Representational Similarity Analysis

We were interested in the representation of the identity of the predictors. To establish whether neural activity carried information about the identity of predictors, we compared the neural activity that we recorded when the same predictors were presented and compared this to situations when different predictors were presented. In both cases the analyses were conducted within our ROIs. To do this, we set up a new whole brain first-level analysis from which we extracted the resulting COPE (contrast of parameter estimate) maps for each session for each ROI. We used the same approach as in the exploratory MRI whole-brain analysis. In other words, we used the same set of regressors and time-locking for the decision and feedback phase as described above (‘Exploratory MRI whole-brain analysis’). However, we changed the way of modelling the confidence phase in order to implement a representational similarity analysis (RSA): for each participant, we categorised data into bundles of three trials of the same predictor and each bundle was coded by a different regressor. Because we reasoned that identity representations should be constructed and strengthened with repeated exposure to the same predictor, the trial bundles were constructed from the end to the beginning of the block. For example, the last three occasions on which a predictor was picked comprised one bundle, the preceding three trials with the same predictor comprised the next bundle and so on. Predictor selections from the very beginning of a block that did not make a bundle of three trials were excluded from the analysis. All trial bundles were submitted to a first-level whole-brain analysis with each bundle represented by a separate regressor. Each regressor consisted of three entries time-locked to the response within the confidence phase as this was the phase of interest for detecting differences in neural representations of social and non-social cues. In other words, each bundle was time-locked to the response made in the confidence phase for the particular trials that were included into the bundle. Duration was set to zero. No other regressors – parametric or non-parametric - were time-locked to these events for the confidence phase. We extracted activation from the same independent dmPFC and bilateral pTPJ ROIs as described above and derived the Euclidean distance for all pairwise combinations of z-maps. We calculated the Exemplar Discriminability Index (EDI) by subtracting the average distances between z-maps of the same predictors from the average distances between bundles of different predictors (while taking care not to include the distance between two instances of precisely the same bundle; Figure 3e). A negative EDI is expected when representations of the same cues are less dissimilar than representations of different cues (Figure 3f). Two participants were excluded as they represented outliers for both brain regions (dmPFC and pTPJ) and for both session (social and non-social). We determined outliers according to three times of the Median Absolute Deviation (matlab function: ‘isoutlier’):

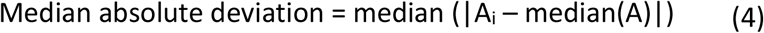

with A representing a variable vector and i=1, 2,…N.

## Acknowledgements

We would like to thank Laurence T Hunt and Geoffrey Bird for helpful comments on task design and analysis. We would like to thank Miriam C Klein-Flügge, Kentaro Miyamoto, Lisa Spiering and Sankalp Garud for helpful comments on earlier versions of the manuscript. The study was funded by a DTC ESRC studentship (ES/J500112/1) to NT and by a Wellcome Trust grant (WT100973AIA) to MFSR.

## Authors contributions

NT, MFSR and MKW designed the experiment. NT collected and analysed the data. NT, MFSR and MKW interpreted the results. MFSR and MKW supervised the project. All authors wrote the manuscript.

## Data availability

We will deposit all choice raw data in an OSF repository. Behavioral results in this paper are derived from these data alone. The access code will be made available.

## Code availability

The above repository will also comprise the full Matlab behavioral analysis pipeline including regression analyses and plotting scripts. A README inside the repository explains the details of its use. The access code will be made available.

