## Supplementary Information for "Neural activity tracking identity and confidence in social information"

**Supplementary figures**

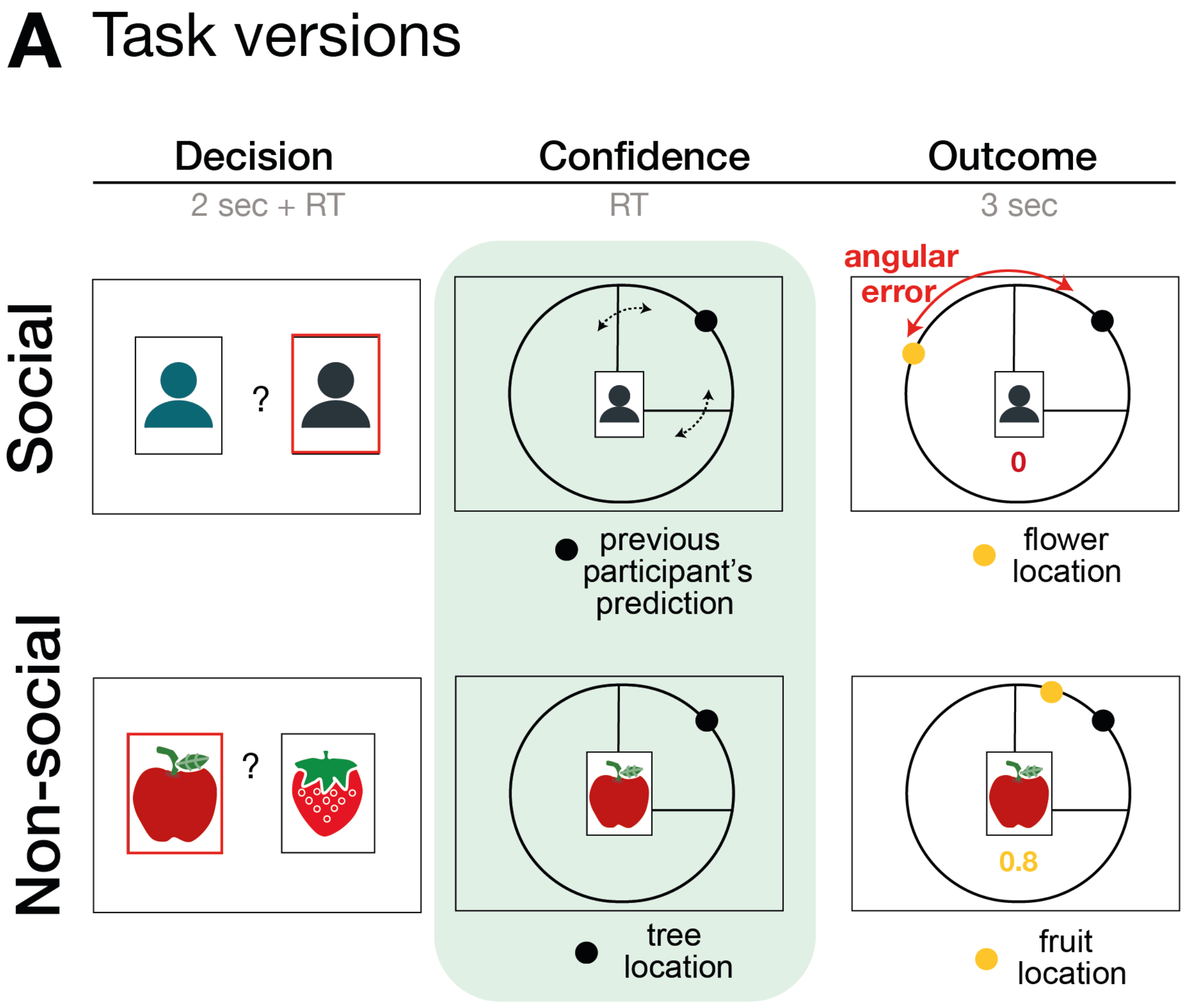


**Supplementary Figure 1. Task design.** The task consisted of three phases: decision, confidence and outcome phase. In the decision phase, participants select between two predictors (either facial stimuli for the social version (Supplementary Table 2, (Lundqvist et al., 1998)) or fruits (Foroni et al., 2013) for the non-social version). Note that here we use abstract person and fruit stimuli, while participants saw respectively real facial and fruit stimuli during the experiment. In the second phase of the trial, participants make a confidence judgment about the performance of the selected predictor. Decisions between predictors did not differ across a variety of behavioural and neural analyses between social and non-social conditions, for details refer to (Trudel et al., 2020). Here we focus on choice and neural signals during the confidence phase.

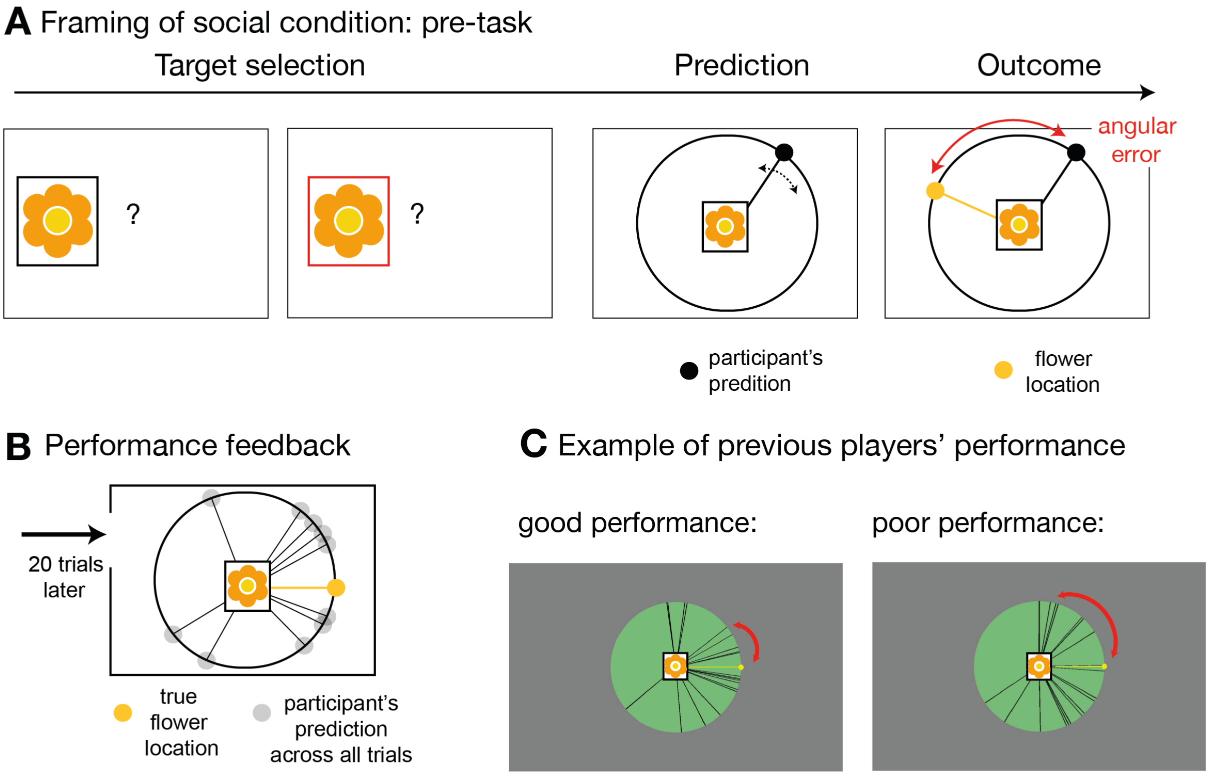


**Supplementary Figure 2. Social instruction procedure.** In the socially framed version of the experiment, participants were instructed that predictors represented previous players who had already performed a different type of behavioural task. In this task, these predictors, or advisors, allegedly had the opportunity to learn about the distribution of a target's location. We will refer to this as pre-task. **(A)** Participants were given explanations of the pre-task performed by the advisors in the following way: on each trial of the pre-task, these other players first selected the target (target selection) and then indicated the location in which they expected the target to appear (prediction). The target was always the same. During the outcome phase, the other players were able to update their beliefs about the target's distribution by comparing the distance between the predicted location and the true location. This distance was defined as angular error. The aim of the advisors was to predict the target as accurately as possible across trials, i.e. to minimize the angular error. In other words, during the pre-task, the other players had learned directly about the target location, while in the main experiment, participants could only infer the target locations from the predictions made by these previous players who now acted advisors, but not directly from the target. Participants in the social condition were instructed that when they picked an advisor to help them predict the target location during the main fMRI experiment, then that advisor would retain the set of trial-by-trial angular errors from the pre-task. As a consequence, in the main task, participants were incentivized to try and identify an advisor who was more accurate in their predictions and they were able to do this by observing the angular errors associated with their predictions (Figure 1A). Note that the main task comprised sets of good and bad predictors and therefore, for the social version, it was important to make it plausible that people might differ considerably in their ability to estimate the true target location. To emphasize the existence of individual differences in advisors' performance, we programmed the pre-task and participants performed 20 trials of the pre-task prior to the main experimental task. The rules were identical to those of the task that participants were told that the advisors had performed. In other words, the participants experienced the pre-task the alleged advisors performed. **(B)** After performing the pre-task themselves, participants were shown their overall performance, which was often normally distributed around the target location, very similar to the prediction distribution observed in the main task. **(C)** In addition, participants were shown similar summary figures of pre-task performance from someone who performed well (left panel) or poorly (right panel) in predicting the target location. Individual differences amongst participants were believable as locating the target on each trial was a challenging task. Importantly, the purpose of the pre-task and subsequent explanation used in the social condition was to make it believable that previous participants, who might now be chosen as advisors in the main task, might have performed well or poorly and therefore might now be more or less accurate predictors in the main task.

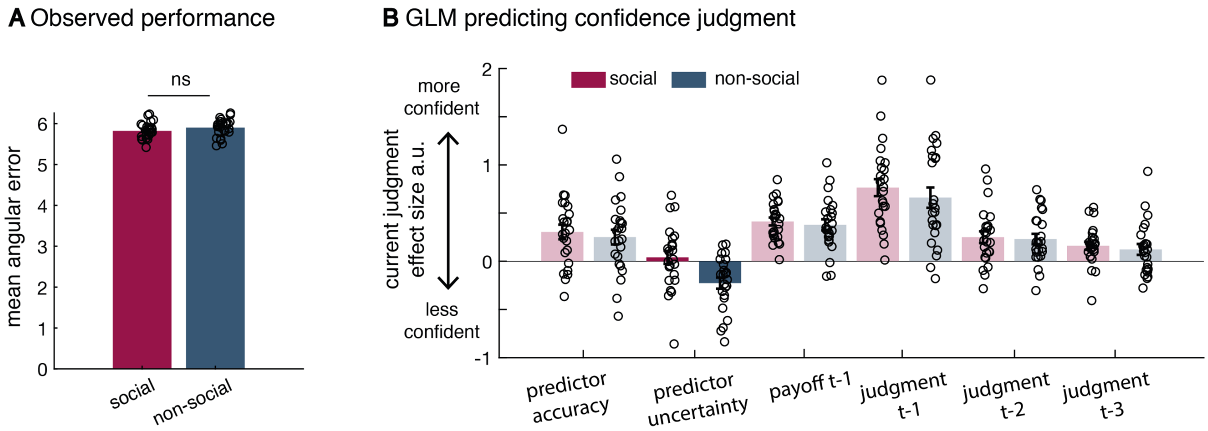


**Supplementary Figure 3. Additional behavioural results. (A)** There was no significant difference in the mean angular error observed between the social and non-social version (mean across angular errors, paired t-test, t(23)=-1.18, p=0.25). There was also no difference in the variance of observed angular errors between conditions (standard deviation across angular errors, Levene test, F(1,46)=0.242, p=0.16; not displayed here). **(B)** All regressors included in the behavioural GLM (t refers to trial); relevant effects are displayed in Figure 2. (n=24, error bars denote s.e.m. across participants.)

Relationship between task parameters and Bayesian belief formation

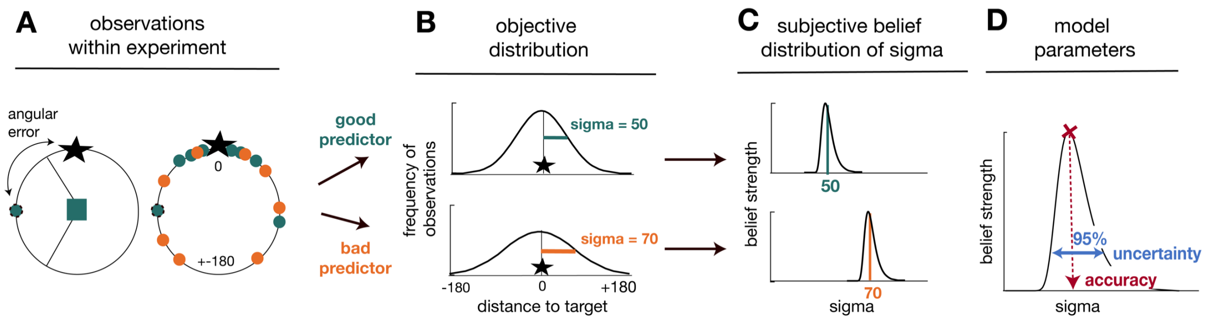


**Supplementary Figure 4. Task statistics and Bayesian model.** Panels depict the mapping between observations during the task (A), their statistical properties (B), and subjective beliefs about these properties derived with Bayes' rule (C,D). **(A)** A predictor's performance can be evaluated by the angular error at each trial (left panel), and by comparing angular errors between predictors across observations (right panel). Better predictors have on average smaller angular errors (green is better than orange). **(B)** Predictors' angular errors were derived from normal distributions centred on the true target location. Critically, the normal distributions for good and bad predictors differed in their standard deviation (sigma): smaller sigma's reflected smaller angular errors, i.e. more accurate predictions of the true target location. Learning about a predictor's angular error across time corresponded to forming beliefs about a predictor's sigma value. **(C)** To capture this learning process, we used Bayesian modelling and derived trial-wise belief distributions over sigma for each predictor. In other words, we estimated a probability density function that expressed the belief strength in each possible sigma over a large range of sigmas, and that was updated with each new observation via Bayes' rule. The coloured vertical lines indicate the true underlying sigmas of the predictors and the black distributions reflect the Bayesian approximation after extensive training. **(D)** We captured two separable estimates about participants' beliefs concerning predictors: an estimate of the accuracy of a predictor (the mode of the distribution indicated by the position of the vertical line on the abscissa) and the uncertainty in that belief (width of the belief distribution). We applied the Bayesian model separately to social and non-social versions. The figure and legend were extracted and adapted from (Trudel et al., 2020).

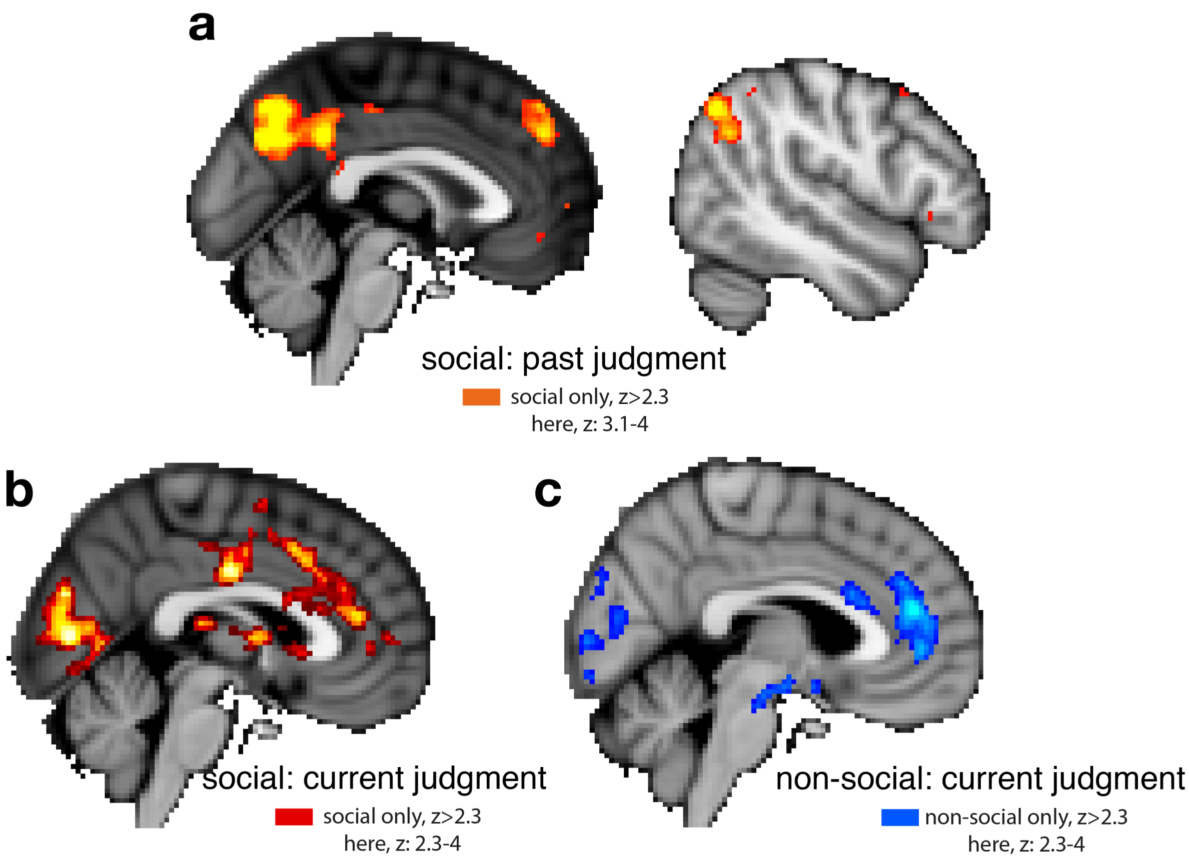


**Supplementary Figure 5. Whole-brain cluster corrected effects for past and current confidence judgments in social and non-social conditions.** For visualisation purposes, brain activations are shown with different z-threshold. All z-thresholds at which we show the activation are denoted under each brain picture. Whole brain-effects are family-wise error corrected with z-score>2.3 and p<0.05. Both regressors, past and current confidence judgments, comprise the selected interval size; hence, a larger interval represents lower confidence in the predictor's performance. To make both regressors more intuitive, we sign-reversed their relationship such that a larger value now also reflects greater confidence in the predictor's performance. Note that we used the same color scheme throughout the report, red indicative for social and blue indicative for non-social condition; please refer to each description for the polarity of each activation. **(a)** Cluster-corrected effects covaried negatively with activation in areas often involved in navigating social scenarios: pTPJ, dmPFC and Precuneus. There were no cluster-corrected effects related to past confidence judgments for the non-social condition and therefore none are shown here. **(b)** Judgments that were made about social advisors in the current trial covaried positively with activation in anterior cingulate cortex with peak activation in ACC, possibly ACC gryus (x/y/z MNI coordinate: 4,40,14) and bilateral striatum (left striatum, x/y/z MNI coordinate: 6, 18, 0). **(c)** Judgments made in the current trial about non-social cues covaried positively with activation in ACC, peaking possibly in ACC sulcus (x/y/z MNI coordinate: 6, 42, 18) (Apps and Sallet, 2017; Apps et al., 2016; Caruana et al., 2018).

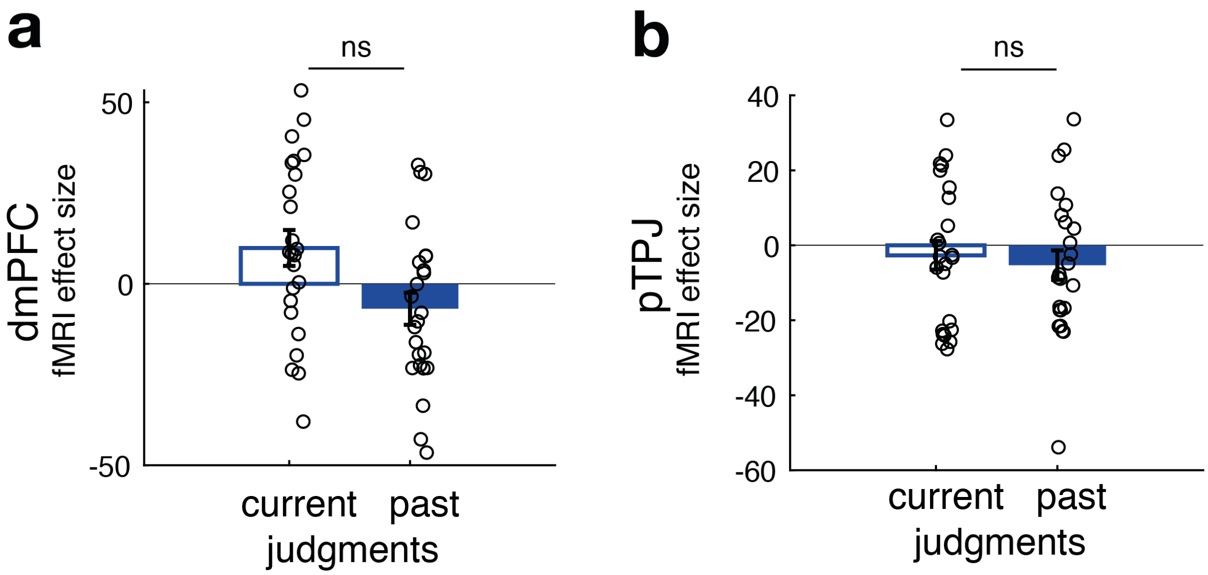


**Supplementary Figure 6. No difference between neural activity of current and past judgments for non-social predictors in dmPFC and pTPJ.** This panel relates to Figure 2e,f and shows that, in contrast to effects found for social predictors, there was no significant difference between the effects of current and past confidence judgments for non-social cues (i.e. the width of the confidence interval set by the participant) on either dmPFC (t(23)=2, p=0.06) or pTPJ (t(23)=0.37, p=0.7) activity. Further, none of the simple effects in either region of interest were significant. (n=24, error bars denote s.e.m. across participants.)

**Supplementary table 1, related to Figure 3 and Supplementary Figure 5.**

| Contrast | Region | Peak Coordinates x/y/z (in mm MNI Space) | Z Value |
|---|---|---|---|
| **Social condition:** past judgment | dmPFC | 2, 46, 34 | -4.1 |
| | Right pTPJ | 48 -64 42 | -4.4 |
| | Inferior Parietal gyrus (left) | -46 -58 48 | -3.8 |
| | Precuneus | -2, -64, 38 | -4.6 |
| | Dorsolateral prefrontal cortex | -46 34 -12 | -3.6 |
| | Left STS | -64 -26 -6 | -3.5 |
| **Social condition:** current judgment | Striatum (right) | 6 18 0 | +3.6 |
| | ACC gyrus | 4 40 14 | +3.6 |
| | Insular (right) | 4 12 0 | +3.8 |
| **Non-social:** current judgment | ACC sulcus | 6 42 18 | +4.4 |
| | Insular (right) | 42 22 0 | +4.1 |
| | OFC (bilateral, here left) | -38 18 -12 | +4.2 |
| Family-wise error cluster corrected, $z > 2.3$, $p < 0.05$ | | | |
